# Spatiotemporal identification of druggable binding sites using deep learning

**DOI:** 10.1101/2020.02.20.952309

**Authors:** Igor Kozlovskii, Petr Popov

## Abstract

Identification of novel protein binding sites expands «druggable genome» and opens new opportunities for drug discovery. Generally, presence or absence of a binding site depends on the three-dimensional conformation of a protein, making binding site identification resemble to object detection problem in computer vision. Here we introduce a computational approach for the large-scale detection of protein binding sites, named BiteNet, that considers protein conformations as the 3D-images, binding sites as the objects on these images to detect, and conformational ensembles of proteins as the 3D-videos to analyze. BiteNet is suitable for spatiotemporal detection of hard-to-spot allosteric binding sites, as we showed for conformation-specific binding site of the epidermal growth factor receptor, oligomer-specific binding site of the ion channel, and binding sites in G protein-coupled receptors. BiteNet outperforms state-of-the-art methods both in terms of accuracy and speed, taking about 1.5 minute to analyze 1000 conformations of a protein with ~ 2000 atoms. BiteNet is available at https://github.com/i-Molecule/bitenet.

## I. INTRODUCTION

Proteins serve biological functionality of a cell via local intermolecular interactions that take place in spatial regions, called binding sites. Binding sites are one of the key elements in drug discovery, being «hot spots» in the pharmacological targets, where the designed drug-like molecule should bind. Identification of novel binding sites expands «druggable genome» and opens new strategies for therapy and drug discovery [1]. Typically drug-like molecules target either orthosteric binding site, where protein interacts with endogenous molecules, or topologically distinct allosteric binding sites [2]. The latter is of a special interest, because allosteric binding sites exhibit higher degree of sequence diversity between protein sub-types, thus, allowing to design more selective ligands, in contrast to the orthosteric ligands [3–5].

Proteins are flexible molecules, that adopt various conformations during their life cycle; and a binding site is a dynamic property of a protein mediated by its conformational changes [6, 7]. Single protein structure represents only minor part of the entire conformational ensemble, hence, binding sites might be easy to overlook from the experimentally determined three-dimensional protein structures [8, 9]. Moreover, many proteins perform their function assembling to oligomeric structure and can form binding sites by means of oligomer’s subunits [10, 11].

Experimental identification of binding sites, such as fragment screening and site-directed tethering [12, 13], using antibodies [14], small molecule microarrays [15], hydrogen-deuterium exchange [16] or site-directed mutagenesis [17] are resource-consuming and may result in negative outcome. On the other hand, computational methods allow to perform large scale binding site identification, investigate protein flexibility via molecular dynamics simulation, and probe to fit chemical compounds using virtual ligand or fragment-based screening.

The classical approaches typically employ empirical scoring functions based on the structural information about known binding sites, or use this information as features for the machine learning algorithms [18–28]. The success rate of these approaches critically depends on the designed features, and may result in false positive predictions, that is identification of «undruggable» regions [29]. Most recently, deep learning approaches, that do not require hand-crafted feature engineering, demonstrated feasibility to predict protein binding sites [30]. In spite of present progress, large-scale binding site detection remains to be a challenge, let alone that there is still a big room for improvement in terms of the method’s accuracy [28].

In this study, we present rapid and accurate deep learning approach, dubbed BiteNet (**Bi**nding si**te**neural **Net**work), suitable for the large-scale and spatiotemporal identification of protein binding sites. Inspired by the computer vision problems, such as object detection in images and videos, we consider protein conformations as the 3D images, binding sites as the objects on these images to detect, and conformational ensembles of proteins as the 3D videos to analyze. We showed that BiteNet is capable to solve the most difficult binding site detection challenges, by applying it to three-dimensional structures of pharmacological targets, including ATP-gated cation channel, epidermal growth factor receptor, and G protein-coupled receptor. Namely, BiteNet correctly identified i) oligomer-specific allosteric binding site formed by the subunits of the trimeric P2X3 receptor complex; ii) conformation-specific allosteric binding site of the epidermal growth factor receptor kinase domain. BiteNet can be used for spatiotemporal investigation of novel binding sites, as we showed by the example of molecular dynamics simulation trajectory for the adenosine A2A receptor. BiteNet outperforms state-of-the-art methods both in terms of accuracy and speed as demonstrated on several benchmarks. It takes approximately 0.1 seconds to analyze single conformation and 1.5 minutes for BiteNet to analyze molecular dynamics trajectory with 1000 frames for protein with ~ 2000 atoms, making it suitable for large-scale spatiotemporal analysis of protein structures. BiteNet is available at https://github.com/i-Molecule/bitenet.

## II. RESULTS

### A. BiteNet architecture

To develop BiteNet we trained 3D convolutional neural network using manually curated protein structures from the Protein Data Bank as the training set (see Section IV). Figure 1 presents the BiteNet workflow. Similarly to 2D images, that have two dimensions (width and height) and three channels for each pixel (red, green, and blue), we represent proteins as 3D images with three dimensions (width, height, and length) and 11 channels for each voxel, where channels correspond to the atomic densities of a certain type (see Section IV) (Figure 1**A**). As neural networks typically take fixed size tensors for the input, we used voxel grid of 64 × 64 × 64 voxels and voxels of 1Å × 1Å × 1Å size. If protein exceeds 64Å in any of the dimensions, we used several voxel grids to represent it (Figure 1**B**). The obtained voxel grids are processed with the 3D convolutional neural network (Figure 1**C**) to output 8 × 8 × 8 × 4 tensor, where the first three dimensions correspond to the cell coordinates relatively to the voxel grid (region of 8 × 8 × 8 voxels), and the four scalars of the last dimension correspond to the probability score of the binding site being in the cell and its Cartesian coordinates. This is followed by the processing of the obtained tensors to output the most relevant predictions of the binding sites (Figure 1**D**). Thus, the input to the BiteNet is the spatial structure of a protein and the output is the centers of the predicted binding sites along with the probability scores. Finally, BiteNet identifies the amino acid residues of a binding site within 6Å neighbourhood with respect to the predicted center. Additionally, when applied to the conformational ensemble of a protein, the obtained predictions and identified amino acid residues are grouped using clustering algorithms (see Section IV A).

**FIG. 1.**
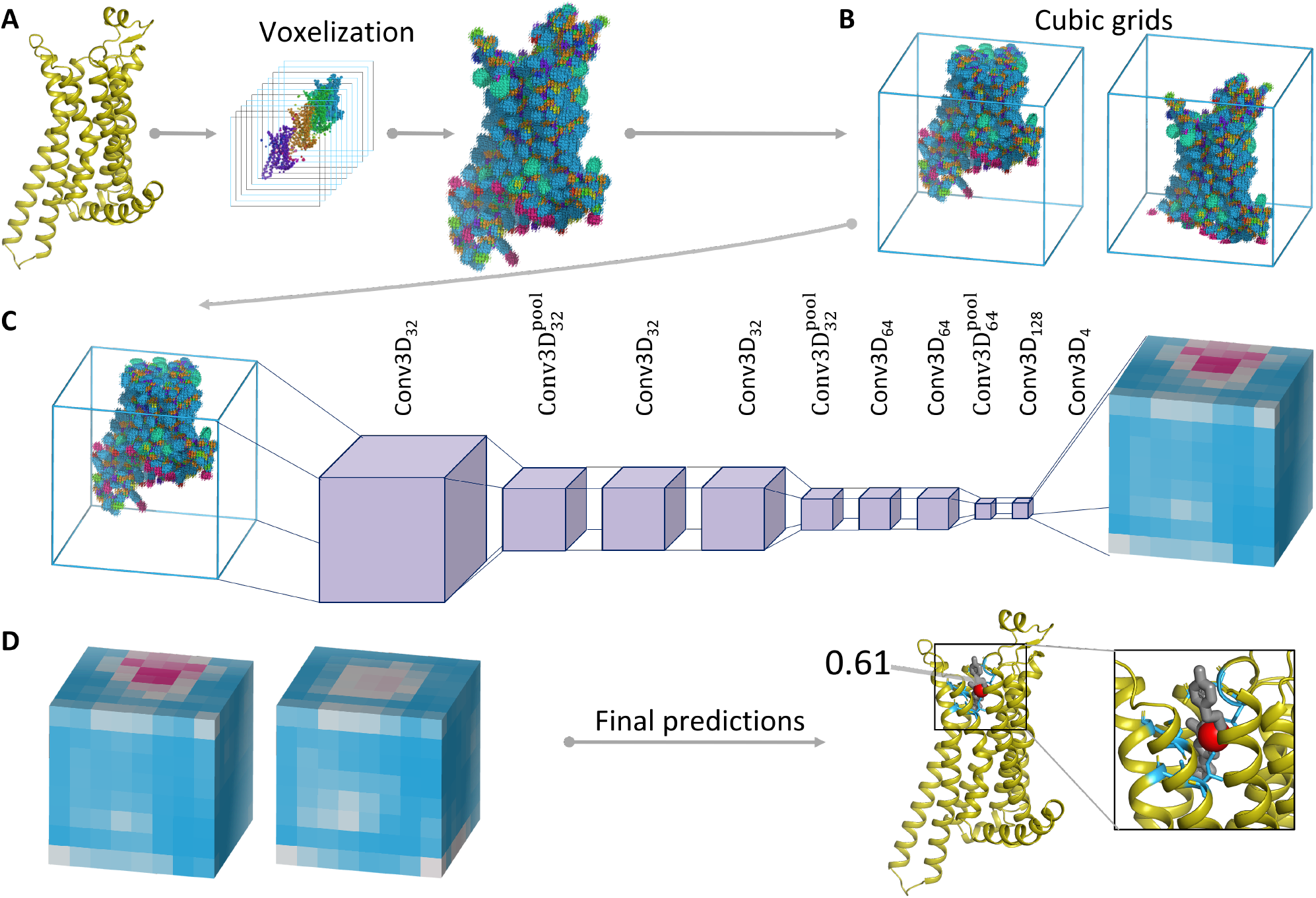
Schematic representation of the BiteNet workflow. (**A**) The input three-dimensional structure of a protein is represented with voxel grid, where channels correspond to the atomic densities. (**B**) The voxel grid is split into fixed-size cubic grids to be fed a neural network. (**C**) Each cubic grid is processed with the 3D convolutional neural network to predict binding sites in fixed-size cells. Cells in cubic grids are colored according to the probability score confidence, from blue to red. (**D**) Predictions obtained for each cubic grid and then processed to output center of the binding site (red sphere), its probability score, and amino acid residues within 6Å of the predicted center (blue sticks). Co-crystallized ligand is shown with gray sticks.

### B. Spatiotemporal prediction of binding sites in pharmacological targets

To demonstrate applicability of BiteNet we considered one of the most difficult binding site detection challenges, comprising three pharmacological targets: the P2X3 receptor of the ATP-gated cation channel family, the epidermal growth factor receptor of the kinase family, and the adenosine A2A receptor of the G-protein coupled receptor family.

#### 1. ATP-gated cation channel

The ATP-gated cation channel, formed by the P2X3 receptor, mediates various physiological processes and represents pharmacological target for hypertension, inflammation, pain perception and others [31]. The channel consists of three identical monomers traversing the membrane, and the orthosteric ATP-binding site comprises amino acid residues of two monomers (see Figure 2**C**). [32]. Drug design targeting the orthosteric binding site is difficult due to highly polarized ATP-specific interface, on the other hand, allosteric ligands targeting protein-protein interactions form promising avenue for drug discovery [11]. Recently allosteric binding site formed by two monomers of a channel was discovered for the P2X3 and P2X7 receptors [11, 33]. We applied BiteNet to the ATP-bound and (AF-219)-bound structures of the trimer complex formed by the P2X3 monomers (PDB IDs : 5SVK, 5YVE), as well as to the single monomer structures. BiteNet correctly identified the orthosteric binding site in the ATP-bound structure and the allosteric binding site in the (AF-219)-bound structure of the trimer, and not in the monomer structures (see Figure 2). Interestingly, BiteNet also predicted center for the ATP-binding site located on the opposite end of the ATP molecule with lower probability score (see Supplementary Figure 1). To ensure, that this is not an artefact of the rotational variance of the model, we generate 50 replicas by rotating the monomer about 10 axes by *π/*3, 2*π/*3*, π,* 4*π/*3, and 5*π/*3 angles and averaged the obtained predictions. As one can see from Figure 2 **D**, although the absolute values of the probability scores vary with respect to the monomers, in all the cases BiteNet correctly identifies the allosteric binding site for the trimer complex and not for the monomer. Note, that ATP is endogenous agonist, while AF-219 is antagonist for the P2X trimer. The agonist-bound and the antagonist-bound conformations are different, particularly, in the regions of the orthosteric and allosteric binding sites (Figure 2 **C,D**). Therefore, BiteNet is sensitive to the conformational changes, as it does not predict the ATP-binding site in the (AF-219)-bound structure and vice versa. Interestingly, despite absence of binding site in the monomer structure, BiteNet predicted different binding sites with relatively high score in the monomer structures. Closer look into available three-dimensional structures of the P2X3 receptors revealed cation ions (Mg, Na, Ca) and ethylene glycol molecules corresponding to these predictions (PDB IDs: 5YVE, 5SVS, 5SVT, 5SVJ, 5SVR, 5SVQ, 5SVP, 5SVM, 5SVL, 6AH4, 6AH5).

**FIG. 2.**
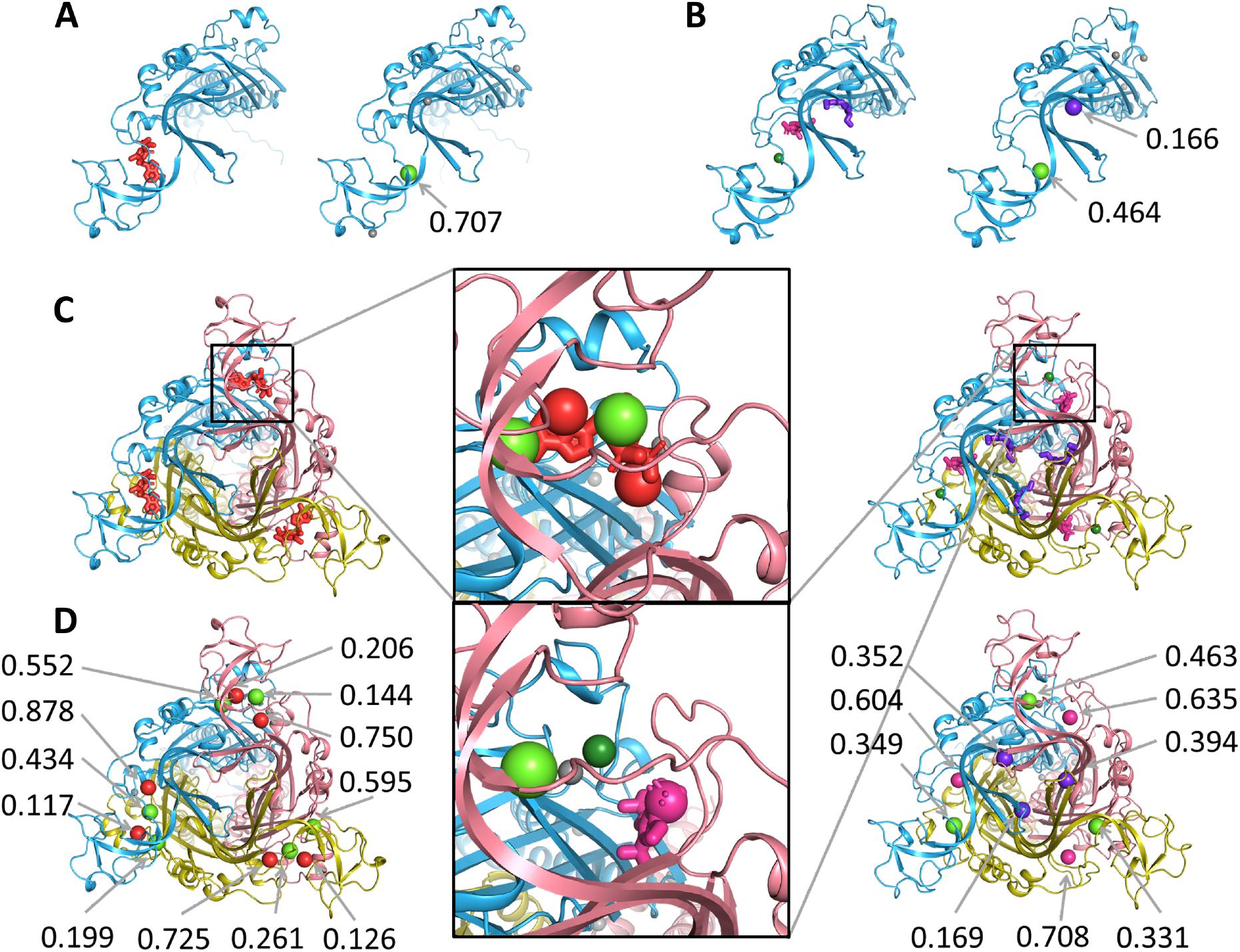
BiteNet predictions for the monomer and oligomer structure of the P2X3 receptor. (**A**) Monomer structure with the orthosteric ligand and cation ion (left), and BiteNet predictions for the monomer structure (right). (**B**) Monomer structure with the allosteric ligand, cation ion and ethylene glycole (left), and BiteNet predictions for this structure (right). (**C**) Agonist-bound (left) and antagonist-bound (right) structures of the P2X3 trimer. (**D**) BiteNet predictions for the agonist-bound (left) and antagonist-bound (right) structures of the P2X3 trimer. Orthosteric and allosteric ligands are shown with red and magenta sticks, respectively. cation ions are shown as dark green spheres and ethylene glycol molecules are shown with violet sticks. BiteNet predictions for these molecules are shown as spheres with the corresponding color.

#### 2. Epidermal growth factor receptor

The epidermal growth factor receptor (EGFR) is a transmembrane protein from the tyrosine kinase family. Over-expression of EGFR is associated with various types of tumors. Although there are EGFR inhibitors targeting the orthosteric binding site of the kinase domain, proteins found in cancer cells often have amino acid substitutions making it insensitive to such inhibitors. There are also mutant-selective irreversible inhibitors that covalently bind to the Cys797 amino acid residue, however, some mutant type receptors possess different amino acid residue at 797 position as well [34]. Recently, three-dimensional structure of L858R/T790M EGFR kinase domain variant bound to the mutant-selective allosteric inhibitor EAI001 was discovered (PDB ID : 5d41) [35]. It was shown, that EAI001 binds to only one monomer, leading to incomplete inhibition, but decreasing cell autophosphorylation. Accordingly, the three-dimensional structure is asymmetric dimer with one monomer bound to both orthosteric and allosteric ligands (the ATP-analogue adenylyl-imidodiphosphate (AMP-PNP) and EAI001, respectively), while the other monomer bound to AMP-PNP only. BiteNet successfully identified both orthosteric and allosteric binding sites in one monomer (chain A) and only former in the other monomer (chain B). We would like to note, that another EGFR kinase domain structure (PDB ID: 5UG9) was in the training set, however, it contains only orthosteric ligand aloof the allosteric binding site.

Although this and previous examples clearly demonstrate BiteNet’s capability to detect binding sites in *holo* conformations, on practice, such conformations can be unknown, especially, when one wants to discover novel binding sites. To evaluate BiteNet’s ability to detect binding sites starting from the unbound conformation, we emulated unbound-to-bound conformational transition as it follows. First, we modelled missing residues in chain B and placed EAI001, as it is observed in chain A. Then, we prepared molecular dynamics system containing chain B, AMP-PNP and EAI001, embedded into the water box with ions using the CHARMM-GUI web server [36]. Next, we run full atom energy minimization of the prepared system until convergence using Gromacs [37], resulting in minimization trajectory consisting of ~ 900 conformations. Finally, we removed ligands, ions, and water and applied BiteNet to each frame of the mini-mization trajectory along with its 50 replicas. Figure 3 shows, that the probability score for the allosteric binding site steadily increases, while the energy of the system is decreasing and the root mean square deviation (RMSD) with respect to the allosteric binding site in the starting (unbound) conformation is increasing. Video 1 demonstrates BiteNet predictions along with the minimization trajectory. Note, that the probability score for the orthosteric binding site remains high during the minimization. Also note, that we used 4Å for the non max suppression distance threshold in order to avoid merging of the predictions for orthosteric and allosteric binding sites during post-processing stage of BiteNet. Therefore, BiteNet can be applied for the large-scale spatiotemporal trajectories in order to detect protein conformations that possess binding sites unseen in the original structure.

**Video 1.**
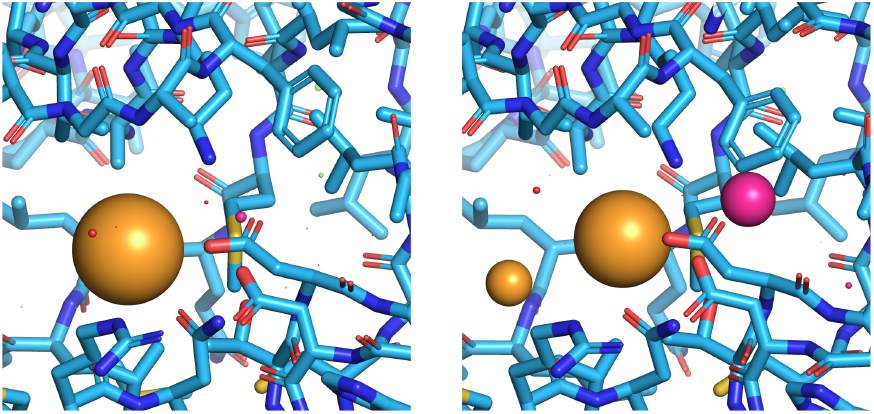
BiteNet applied to the minimization trajectory of EGFR kinase domain starting from the unbound state. Predictions corresponding to the orthosteric and allosteric sites are shown as yellow and magenta spheres, respectively. Frames 1 (left) and 894 (right) are shown.

**FIG. 3.**
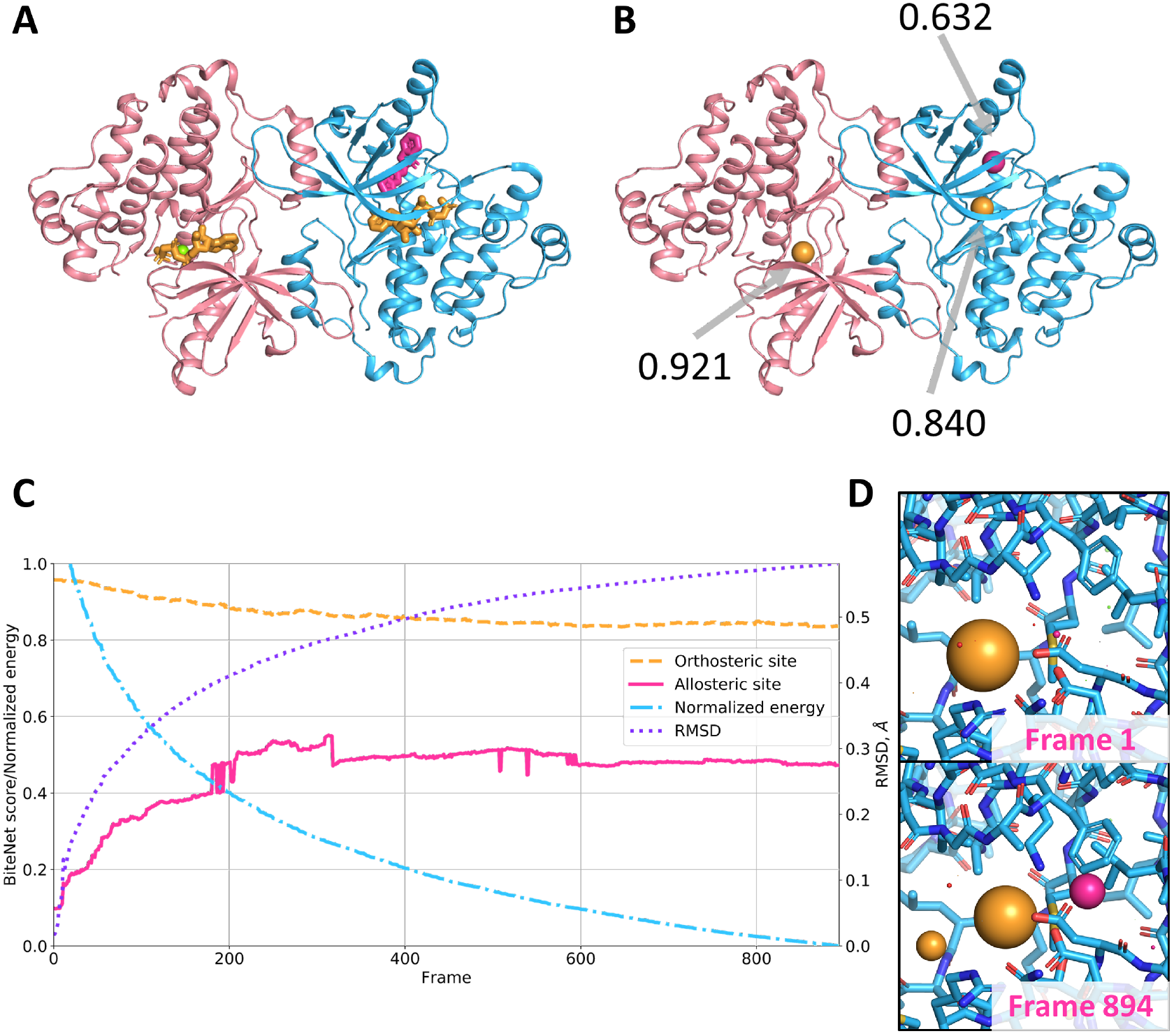
(**A**) Assymetric dimer structure of the EGFR kinase domain. Orthosteric and allosteric ligands are shown with yellow and magenta sticks, respectively, Mg ion is shown as green sphere. (**B**) BiteNet predictions for the assymetric dimer, the predicted centers for the ligands are shown as spheres with the corresponding color. (**C**) BiteNet predictions obtained for the energy minimization trajectory. The normalized energy is shown with blue dash-dotted line, the RMSD with respect to the unbound conformation of the alloteric binding site is shown with violet dotted line, BiteNet probability score for the orthosteric and allosteric binding sites are shown with dashed orange and magenta solid lines, respectively. The normalized energy of 1 and 0 corresponds to −7.76969e+5kJ/mol and 8.80655e+5kJ/mol, respectively. (**D**) The starting and the final conformations of the minimization trajectory along with BiteNet predictions.

#### 3. G protein-coupled receptor

G protein-coupled receptors (GPCRs) mediate numerous physiological processes in the body, making them important targets for modern drug discovery. Most of FDA-approved drugs bind to orthosteric binding sites of GPCRs. However, such drugs may be non-selective with respect to the highly homologous receptor sub-types. In such cases, there is need in drug design targeting allosteric binding sites, that are less conserved than orthosteric one [38]. Three-dimensional structures of GPCRs reveal allosteric binding sites spanning extra-cellular, transmembrane, and intracellular regions; identification of novel allosteric sites in GPCRs can provide alternative options for drug discovery [39]. To demonstrate the use of BiteNet in spatiotemporal identification of GPCR binding sites we analyzed molecular dynamics trajectories of the human adenosine A2A receptor (A2A) retrieved from the GPCRmd repository [40].

Namely, we considered trajectories of A2A embedded into the POPC lipid bilayer surrounded by water, sodium and chloride ion molecules starting from the active-like conformation (PDB ID: 5G53) in complex with agonist NECA and with no ligand (GPCRMD IDs: 48:10498 and 47:10488, respectively). In total each simulation lasted for 500*ns* with the time step of 4.0*fs* and interval between frames of 2.0*ns*, resulting in 2500 conformations of A2A. We consequently applied BiteNet for each frame of the trajectory. As expected, in both simulation trajectories we observed a cluster of predictions corresponding to the canonical orthosteric binding site in GPCRs. The cluster is more dense and with higher averaged score in the ligand-bound simulation trajectory, which could be explained by lower flexibility of the protein due to the protein-ligand interactions. Surprisingly, in both simulation trajectories we also observed cluster of predictions in the neighbourhood of the end of TM1, TM7 and helix 8 starting from ~300ns in the ligand-free simulation and from ~150 to ~200 ns and from ~320 to ~370 ns in the ligand-bound simulation. Closer look to the conformations with the highest probability scores corresponding to this cluster revealed lipid tail buried to the cavity formed by hydrophobic amino acid residues. It is important to note, that although GPCRs are tightly surrounded by lipids, BiteNet did not produced predictions all over the region exposed to a membrane, as it was explicitly trained on druggable binding sites. To investigate if the lipid tail binds to the cavity, for each frame *f* we calculated its mobility in terms of RMSD between the conformation of the lipid tail in this frame and the conformation of the lipid tail averaged over [*f* − 100*, f* + 100] frames. As one can see from Figure 4, the calculated RMSD is lower for the frames with high probability scores corresponding to the predicted binding site. Videos 2 and 3 demonstrates BiteNet predictions and binding of the POPC molecule during these simulations. To the best of our knowledge there is no available structures for any GPCR with ligand bound to this region. When applied BiteNet to molecular dynamics trajectories obtained for other receptors from GPCRmd, we also observed similar cluster in the muscarinic M2 receptor, again, starting from active-like conformation (data not shown). Thus, the predicted region may be worth paying attention to, as it may correspond to the novel allosteric binding site in GPCRs.

**Video 2.**
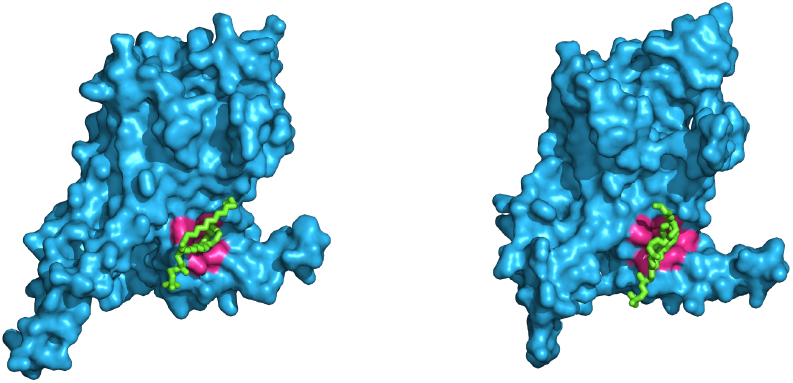
BiteNet applied to the ligand-free A2A simulation trajectory. BiteNet predictions for the orthosteric and hypothethical binding sites are colored with yellow and magenta, respectively. Frames 1489 (left) and 2055 (right) are shown.

**Video 3.**
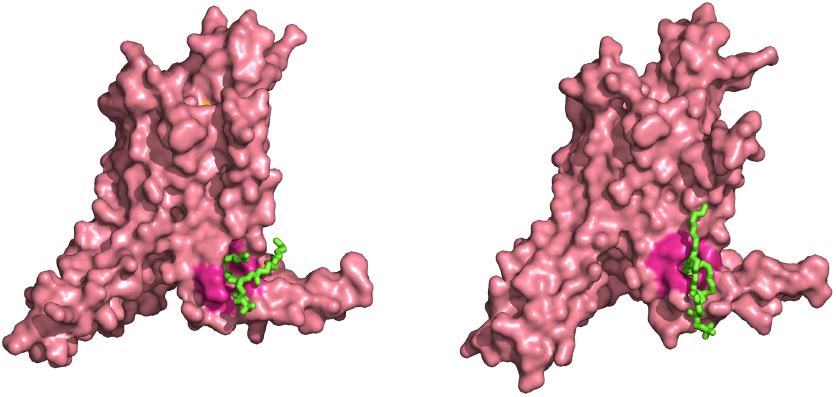
BiteNet applied to the ligand-bound A2A simulation trajectory. BiteNet predictions for the orthosteric and hypothethical binding sites are colored with yellow and magenta, respectively. Frames 835 (left) and 1806 (right) are shown.

**FIG. 4.**
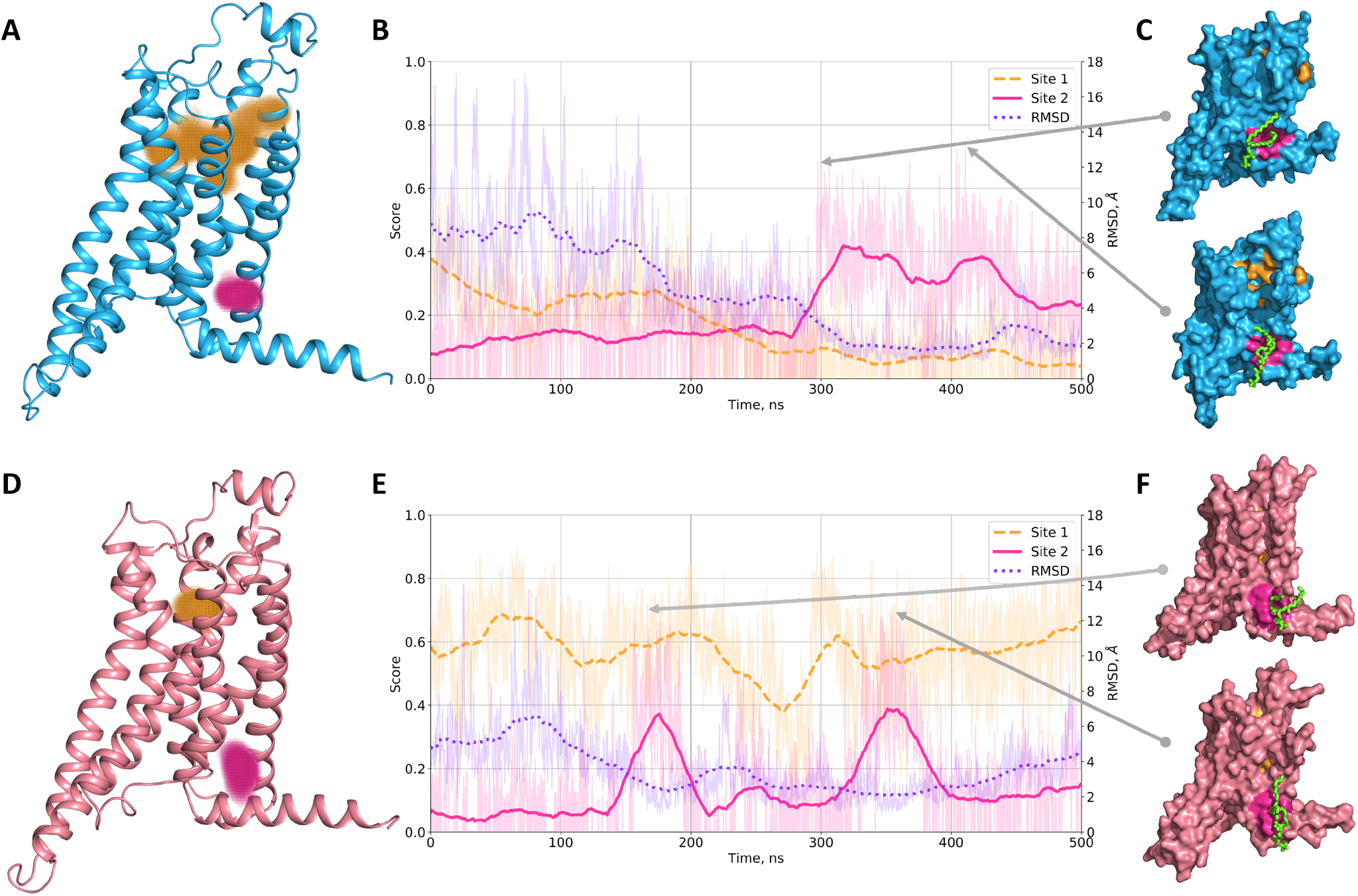
BiteNet predictions for molecular dynamics simulation of the adenosine A2A receptor. (**A**, **D**) Starting ligand-free and agonist-bound conformations of A2A, respectively. Orange point clouds corresponds to the BiteNet predictions of the canonical orthosteric binding site in A2A, while magenta point cloud corresponds to the BiteNet predictions of the hypothetical binding site, observed during the simulation. (**B**, **E**) BiteNet probability scores for the orthosteric binding site (dashed orange line), allosteric binding site (magenta solid line), and RMSD with respect to the window-based mean lipid tail conformation (dotted violet line), computed for the molecular dynamics trajectories. (**C**, **F**) A2A conformations corresponding to the highest BiteNet probability scores for the hypothetical binding site. Lipid tail binding to the hypothetical binding site is shown with green sticks.

To summarize, we showed applicability of BiteNet for binding site detection for three different pharmacological targets and challenging binding sites observed in soluble as well as in transmembrane protein domains. BiteNet was capable to detect conformation-specific and oligomer-specific allosteric binding sites and can be applied for large-scale spatiotemporal analysis of protein structures. Using the example of A2A we demonstrated how BiteNet can be used on practice to investigate novel binding sites. We also would like to note, that used three-dimensional structures were not exposed to BiteNet during the training process. In the next section we demonstrate computational efficiency of BiteNet in terms of accuracy and speed by comparing it against the existing computational methods on binding site prediction benchmarks.

### C. Computational efficiency of BiteNet

To compare BiteNet with the other approaches we evaluated its performance on the COACH420 [41] and HOLO4K [42] datasets, that contain 420 and 4542 proteins, respectively. Note, that for fair comparison we considered only proteins not presented in the method’s train sets, and for which all methods successfully predict true binding sites according to the P2Rank criterion [28], resulting in the 230 and 2305 protein subsets from COACH420 and HOLO4K, respectively.

As the performance metric we calculated the average precision (AP), that is the area below precision-recall curve, for *All* and *TopN* predictions, where *N* is the number of the true binding sites present in a protein structure. As one can see from Figure 5 BiteNet out-performs classical binding site prediction methods, such as fpocket [23], SiteHound [21], MetaPocket [24], as well as the state-of-the-art machine learning methods, such as DeepSite [30] and P2Rank [28] (Supplementary Table 1 lists more detailed comparison including precision and recall metrics).

**FIG. 5.**
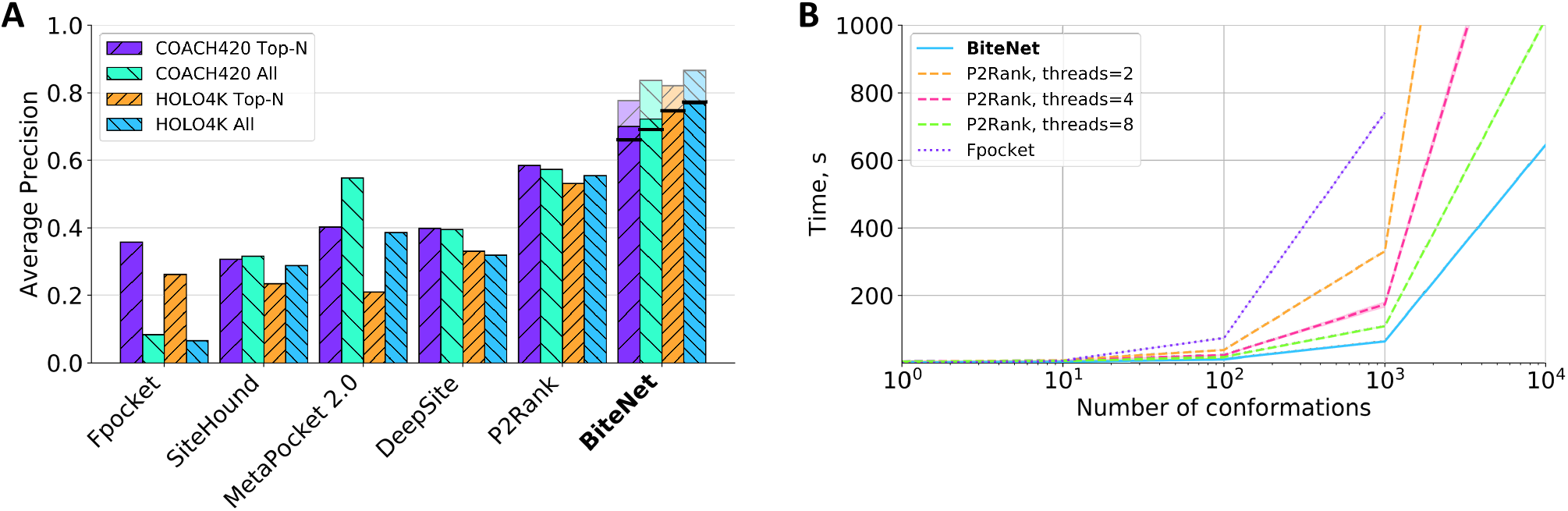
(**A**) Performance of the binding site prediction methods on the COACH420 and HOLO4K benchmarks. *All* and *Top - N* correspond to the average precision calculated taking into account all and the top *N* predictions, where *N* is the number of true binding sites in a protein. Pale bars correspond to the BiteNet performance, when the true positive binding site is defined as in the training. Black lines correspond to the BiteNet performance on the whole benchmarks. (**B**) Elapsed time for fpocket, P2Rank, and BiteNet to analyze 1, 10, 1000 and 10000 conformations of a protein with ~2000 atoms. The computed elapsed time is the average of 10 independent runs.

BiteNet is also computationally efficient, Figure 5B shows elapsed time spent by BiteNet along with fpocket and P2Rank, which are one of the fastest methods, with respect to the number of the processed protein conformations. BiteNet, that runs on a single GPU (GeForce GTX 1080 Ti), outperforms P2Rank that runs on several CPUs (Intel(R) Core(TM) i7-8700K CPU @ 3.70GHz). On average, BiteNet takes approximately 0.1 seconds to process single protein conformation. Further optimization of CPU-GPU interconnection and multiple GPUs implementation of BiteNet will result in even faster performance.

## III. DISCUSSION

In this study we introduced BiteNet, a deep learning approach for spatiotemporal identification of binding sites. BiteNet takes advantages of the computer vision methods for object detection, by representing three-dimensional structure of a protein as a 3D image with channels corresponding to the atomic densities. BiteNet goes beyond classical problem of binding site prediction in *holo* protein structures, exploring protein dynamics and flexibility by means of large-scale analysis of conformational ensembles. The detected conformations with observed binding site of interest, then can be used for structure-based drug design approaches, such molecular docking and virtual ligand screening, as well as structure-based de novo drug design.

We believe superior performance of BiteNet with respect to the other machine learning methods for binding site prediction was achieved due to careful preparation of the training set and training process; below we address several important issues related to these procedures.

### A. Training set

Curated and well-balanced training set is of crucial importance for derivation of machine learning models and its applicability domain. Experimentally determined protein structures often contain detergent and buffer molecules, that reveal electron density. This should be considered carefully and not mixed up with the true binding sites. To avoid potential bias related to this problem we filtered out typical detergent and buffer molecules (see Supplementary Table 2). Note, however, this procedure likely resulted in removing both false and true positives binding sites. For example, we discarded lipid molecules surrounding membrane proteins, including functional lipid molecules, such as cholesterol. Additionally, training set inevitably contains false negative binding sites, because protein structures may also contain empty binding sites. Another source for false positive binding sites come from symmetrical oligomer structures, as for example the P2X3 trimer. Indeed, the asymmetric unit does contain the ligand, however the binding site is formed not only by the asymmetric unit, but also by symmetry mates, which are usually omitted in the analysis. We also observed structures with missing atoms and residues in the binding sites; we believe such structures should be either properly refined or discarded from the training set. In addition, the definition of the true positive prediction and binding site itself may vary. Binding site is typically defined with respect to the cutoff distance between the protein and ligand atoms (4.0Å in this study), center of the binding site can be defined as the center of mass of the ligand or the binding site residues (in this study), and the true positive prediction can be defined with respect to the cutoff distance between the ligand or center of the binding site (4.0Å in this study). We choose the later definition of the true positive prediction because it is invariant with respect to the type of the ligand and its binding pose.

Training-test split is another important issue, that affects performance of the derived model. First of all structural similarity should be taken into account, as it is known that proteins with low sequence similarity may still share highly similar protein fold. We observed that the largest cluster contains 4044 protein chains of similar structures. Splitting this cluster into the train and test sets would likely result in the bias and overfit with respect to the corresponding protein fold. To circumvent this issue we carefully distributed protein structures, such that there is no highly similar structures in the training and test sets in terms of the TM-score structural similarity [43].

Data augmentation techniques can be also helpful to derive more robust predictive models. For protein binding site prediction problem, computational methods to generate conformational ensembles can be used in order to represent binding site with multiple orientations or even small perturbations. In this study, due to computational limitations, we used implicit data augmentation and provided random orientation of proteins to the neural network each epoch.

### B. Training model

Hyperparameters, such as neural network architecture, type of the activation functions, the learning rate, and many others, influences the model performance. Thus, fine-tuning is needed in order to find optimal set of the hyperparameters. We trained several models and found the following hyperparameters to be optimal: 64 voxels for the cubic grid size, 1.0Å for the voxel size, 4.0Å for the density cutoff, 48 for the stride parameter, 16 for the minibatch size, 1e-5 and 10.0 for the *γ* and *λ* parameters, respectively (see Supplementary Table 3 for evaluation of models corresponding to different parameters). Among these parameters, the voxel size has dramatic influence on the computational speed, it takes ~2 times more to train and apply the model with the voxel size of 0.8Å, as compared to the voxel size of 1.0Å. On the other hand, we observed model corresponding to the voxel size of 2.0Å to be faster, though less accurate. Although we achieved satisfied performance of the resulting model (the average precision was improved from 0.4 to 0.53), our parameter screen is not meticulous. The auto-ml approaches would be useful to find optimal model through extensive search of neural network architecture and parameters [44, 45].

Note, that the obtained model is not rotation-translation invariant by construction; it could be easily seen from the different binding site scores assigned to identical subunits of oligomer (see Supplementary Figure 2). To make sure this does not significantly affect BiteNet’s performance, we re-evaluate the average precision on the augmented test set, that contains additional 50 replicas of each protein obtained with rotation by *π/*3, 2*π/*3*, π,* 4*π/*3, and 5*π/*3 angles about 10 different axes corresponding to the centroids of the icosahedron facets [46]. Indeed, we observed that the average precision computed for the *TopN* predictions almost did not change, and became slightly higher when considering *All* predictions. This is because for some replicas BiteNet produced additional binding site predictions with very low scores. Thus, it might be useful to apply BiteNet for different orientations of the structure and average the obtained results.

## IV. METHODS

### Training dataset

To compose the training set we retrieved atomic structures of protein-ligand complexes with resolution better than 3.0Å, that contain less than 4 protein chains, and the sequence identity threshold of 90% from Protein Data Bank (PDB)[47]. Then we refined each protein structure by replacing non-standard amino acid residues with the standard ones, modelling missing residues and short loops (less than 10 amino acid residues) using the ICM-Pro software (molsoft.com). Note, that we did not model N- and C-terminus, as well as long missing loops of more than 10 amino acid residues. Then we discarded proteins, if refinement affects 3 or more atoms of its binding sites, because such conformational changes could be incompatible with the ligand binding pose. We also discarded water molecules, ions, protein chains with length less than 50 amino acid residues, and considered only non-detergent molecules (see Supplementary Table 2) with more than 14 heavy atoms as the ligands. We further disregarded protein complexes with less than 20 protein heavy atoms in the binding site, that is protein atoms within 4Å distance from the ligand. Finally, we manually filtered out “long” proteins, which length across at least one of the principal axis was more than 250Å (see Supplementary Figure 3). This procedure yielded the final set of 5946 atomic structures of protein-ligand complexes comprising 11301 polypeptide chains and 11949 binding sites.

We considered each protein of a protein complex as a voxel grid, with voxel size of 1.0Å with no spacing between the voxels. We represented each voxel by 11 channels corresponding to the atomic density function of acertain atom type, similarly to [48]:

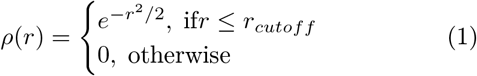

where *r_cutoff_* is the distance threshold of 4Å.

For rigorous validation of the prediction model it is important to carefully split the training and the test datasets. Given that proteins with low sequence similarity may still have high structural similarity, the standard random split would likely lead to the biased training and test sets. To reduce possible bias, we calculated structural similarity for each pair of protein chains in the dataset using the TMalign software [43], resulting in 11301 × 11301 structural similarity matrix (see Supplementary Figure 4). Then we grouped protein chains using the hierarchical clustering algorithm implemented in *sklearn* [49, 50], such that structural similarity of any two protein chains from different clusters is less than 0.5. Finally, we split the dataset in a way that the training and the test sets do not share protein chains from the same clusters, comprising 9844 and 1457 protein chains, respectively.

### Neural network architecture

Given *N_x_ × N_y_ × N_z_ × N_c_* voxel grid representation of a protein, we first divided it into the cubic grids of the fixed shape of 64 × 64 × 64 voxels with stride of 48 voxels, in order to get constant size input for the neural network. We considered cubic grids with the average atom density less than 1e-4 as empty cubic grids, and discarded it from the training and test sets. Following the Yolo approach for the object detection problem in images [51], we constructed neural network that converts 64 × 64 × 64 cubic grid into 8 × 8 × 8 cubic cells of size 8 × 8 × 8 voxels each, and aims to identify target cells, that contain centers of the binding sites, along with the center’s coordinates. Thus, the output of the prediction model is 8 × 8 × 8 × 4 tensor, where the first three dimensions are the cell coordinates with respect to the cubic grid (*i_cell_, j_cell_, k_cell_*), and the four scalars of the fourth dimension are the probability score 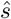 that the corresponding cell contains center of a binding site, and the coordinates of this center with respect to the cell 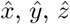. The core of the neural network comprises ten 3D convolutional layers: 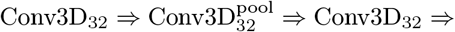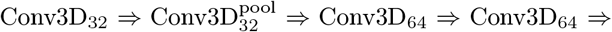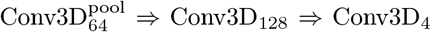, where the sub-script number denotes the number of filters. We used kernels of size (3, 3, 3) for each layer, stride of 2 for the pooling layers, and the batch normalization and the rectified linear unit (ReLu) activation function for all layers, except for the last one. Finally, we use the sigmoid activation function to obtain probability score 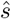 in the range of (0, 1) and relative coordinates 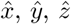 of the predicted center of the binding site with respect to the cell. The Cartesian coordinates are then calculated according to 2:

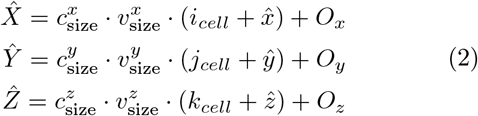

where *c*_size_ and *v*_size_ corresponds to the size of a cell and voxel, respectively, and *O_x_*,*O_y_*,*O_z_* are the Cartesian co-ordinates of the origin of the cubic grid.

We used custom loss function for training, that contains three terms:

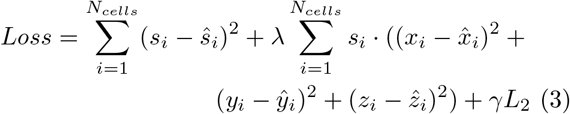

where *N*_*cells*_ is the number of cells in the single cubic grid, s*_i_* and 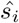 are the true (0 or 1) and predicted probability scores of the cell, *x_i_, y_i_, z_i_* and 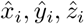 are the true and predicted coordinates for *i*-th cell, respectively, and *L*_2_ correspond to the regularization term. Therefore, the first and the second terms aim to penalize incorrect prediction of the probability score and the center of the binding site, respectively. Note, that we multiply the second term by the true probability score (0 or 1) to take into account only relevant predictions. The third term is the *L*_2_ regularization term for the neural network parameters. The coefficients *λ* = 5 and *γ* = 1e-5 are the weights of the penalty terms.

We trained the network in Tensorflow v1.14 [52] for 400 epochs using the Adam optimizer with the default parameters, minibatch size of 16 cubic grids, and the learning rate of 1e-3 gradually decreasing to 1e-5 during the training. We would like to note, that presented architecture is not invariant to rotations of a protein. Data augmentation, i.e. considering different orientations of a protein within a single epoch, may circumvent this problem to some extent. Because of GPU memory limitations, in this study we used implicit data augmentation by considering random orientation of a protein each epoch.

To obtain the final predictions we applied the post-processing procedure, as it follows. First, we discarded all the predictions with the probability score 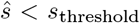. The remaining predictions are then processed by means of the non-maximum suppression. More precisely, we select the best prediction in terms of the probability score, as the seed of a cluster, and put all the predictions with the centers of the binding site closer than *d*_threshold_ = 8Å to the center of the best prediction. Then we select the second best prediction, as the seed of the next cluster, and repeat the above procedure until all the predictions are clustered. Finally, we keep only *N*_top_ seeds in terms of the probability scores, as the final predictions. For the training we used *s*_threshold_ = 0.1 and *N*_top_ = 5, for benchmarking *s*_threshold_ = 0.01 (in order to calculate AP for all predictions), and for the case study *s*_threshold_ = 0.1 and all predictions. To evaluate the performance of the prediction model we define the true positive (*TP*) prediction of the binding site, as the top-scored correct prediction, that is prediction with the probability score 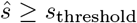 and the predicted center of the binding site within *d*_threshold_ from the true center of the binding site. The rest of the predictions are considered as false positives (*FP*). Given this, we calculate precision and recall metrics according to:

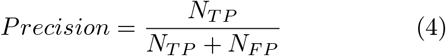

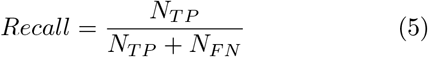

where *N_F N_* is the number of false negative predictions, that is the number of binding sites with no correct prediction. As the main metric we calculate the average precision metric *AP*, which is the area under precision recall curve.

Note, that we define correct prediction with respect to the center of the binding site, rather than binding pose of a ligand. We believe this is more rigorous metric, because it does not depend neither on the binding pose of a ligand, nor on the ligand itself. However, for fair comparison with the existing methods, we also computed the metrics, where the prediction is considered to be true positive prediction, if the minimal distance to the ligand is less than *d*_threshold_ = 4Å.

#### A. Clusterization

Given conformational ensemble of a protein, as for example, molecular dynamics trajectory, we firstly applied BiteNet to each conformation. Then we grouped the obtained predictions using clustering algorithms. In this study we used three different clustering approaches implemented in the sklearn python library [50] : the mean shift clustering algorithm (MSCA) [53], the density-based clustering algorithm (DBSCAN) [54, 55], and the agglomerative hierachical clustering algorithm [49]. While the first two approaches are mainly applied for the set of points in Euclidean space, the latter approach can be applied also for set of amino acid residues forming the predicted binding site. Finally we assigned two scores for each cluster. The first score is the sum of maximal probability score of a cluster in each frame averaged over the total number of frames. For the second score, the mean sum of probabilities scores (larger than *cluster*_*score*_*threshold*_*step* = 0.1) of a cluster is computed for each frame; these sums are then averaged over the total number of the corresponding frames. We implement several clustering approaches, because it is known that clustering results may strongly vary depending on clustering algorithm and different parameters for them, also affecting the cluster scores.

## V. CONCLUSIONS

In this study we introduced BiteNet, a deep learning approach for spatiotemporal identification of binding sites. BiteNet takes advantages of the computer vision methods for object detection, by representing three-dimensional structure of a protein as a 3D image with channels corresponding to the atomic densities. BiteNet goes beyond classical problem of binding site prediction in *holo* protein structures, exploring protein dynamics and flexibility by means of large-scale analysis of conformational ensembles. It is able to detect allosteric binding sites for both soluble and transmembrane protein domains and outperforms state-of-the-art methods both in terms of accuracy and speed. BiteNet takes approximately 0.1 seconds to analyze single conformation and 1.5 minutes to analyze molecular dynamics trajectory with 1000 frames for protein with ~ 2000 atoms. BiteNet is available at https://github.com/i-Molecule/bitenet.

## ACKNOWLEDGMENTS

We acknowledge the HPC team at CDISE (Skoltech) for support usage of the “Zhores” supercomputer in order to train BiteNet.

