## Supplementary Information for "Spatiotemporal identification of druggable binding sites using deep learning"

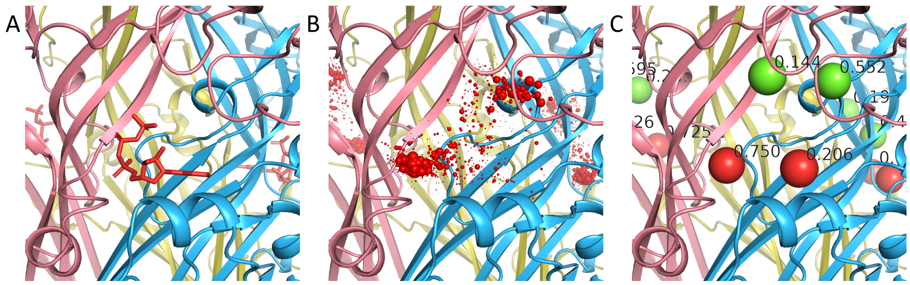


Figure 1: BiteNet predictions for the ATP-bound P2X3 receptor trimer augmented with 50 replicas. (**A**) the ATP-binding site, ATP is shown with red sticks) (**B**) BiteNet predictions with no non max suppression. (**C**) BiteNet predictions filtered by the non max suppression with $distance_threshold = 8$Å.

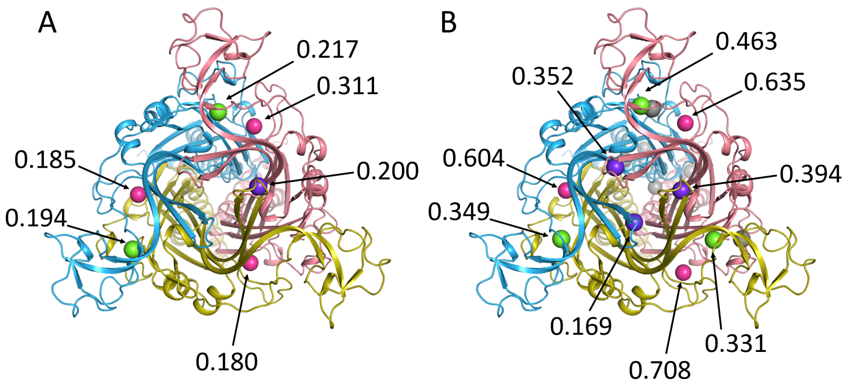


Figure 2: BiteNet predictions obtained for the P2X3 trimer with no (**A**) or with (**B**) rotational replicas.

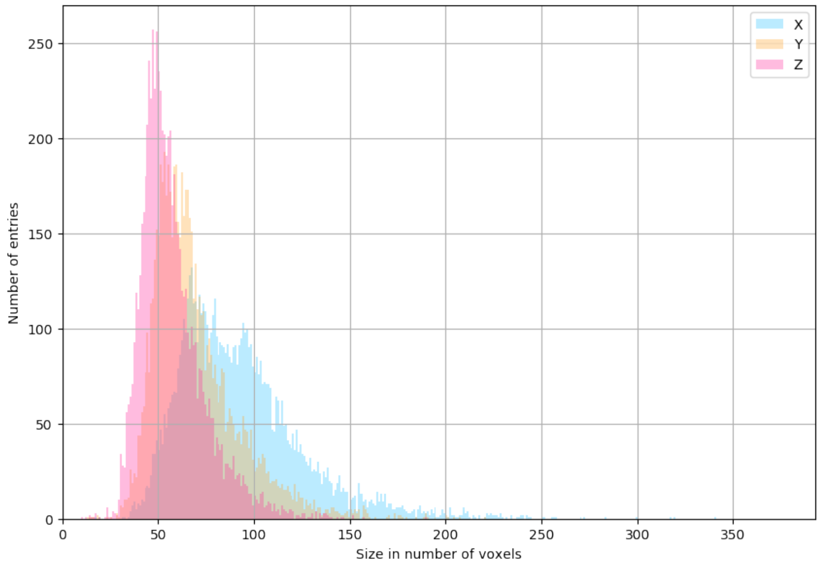


Figure 3: Statistics for the protein lengths measured in voxels with respect to its principal axes.

Table 1: Binding site prediction methods performance on the COACH420 and HOLO4K benchmarks. We calculate three performance metrics, precision, recall and average precision, for top-1, top-3, top-n, top-(n+2) and all predictions, where $n$ is the number of true binding sites in a protein structure. Note that precision and recall metrics may be not indicative, because precision would show higher scores for methods that predict small number or no binding sites, and recall assigns higher score to methods which predict large number of binding sites even with low precision. Average precision metrics doesn't have such disadvantages. BiteNet+ corresponds to BiteNet performance on the augmented dataset, that is with 50 additional rotational replicas. Bold font correspond to the best metric.

| Method | Top-1 | | | Top-3 | | | Top-n | | | Top-(n+2) | | | All | | |
|---|---|---|---|---|---|---|---|---|---|---|---|---|---|---|---|
| | prec | rec | ap | prec | rec | ap | prec | rec | ap | prec | rec | ap | prec | rec | ap |
| Performance on COACH420 dataset based on all protein entries | | | | | | | | | | | | | | | |
| Fpocket | 0.648 | 0.490 | 0.339 | 0.315 | 0.634 | 0.215 | 0.519 | 0.555 | 0.321 | 0.292 | 0.663 | 0.211 | 0.086 | 0.755 | 0.074 |
| SiteHound | 0.585 | 0.439 | 0.289 | 0.285 | 0.641 | 0.329 | 0.469 | 0.531 | 0.329 | 0.257 | 0.668 | 0.333 | 0.020 | **0.933** | 0.338 |
| MetaPocket 2.0 | 0.714 | 0.556 | 0.393 | **0.591** | 0.602 | 0.373 | 0.656 | 0.587 | 0.386 | **0.559** | 0.625 | 0.373 | 0.048 | 0.811 | 0.510 |
| DeepSite | 0.658 | 0.486 | 0.372 | 0.308 | 0.572 | 0.377 | 0.540 | 0.517 | 0.387 | 0.295 | 0.575 | 0.378 | **0.256** | 0.577 | 0.378 |
| P2Rank | 0.810 | 0.623 | 0.536 | 0.332 | 0.707 | 0.552 | 0.642 | 0.666 | 0.557 | 0.309 | 0.723 | 0.557 | 0.152 | 0.740 | 0.553 |
| BiteNet | **0.853** | **0.651** | **0.609** | 0.384 | 0.771 | 0.681 | **0.695** | 0.721 | 0.661 | 0.365 | 0.788 | 0.687 | 0.236 | 0.807 | 0.691 |
| BiteNet+ | 0.851 | 0.646 | 0.609 | 0.345 | **0.784** | **0.698** | 0.663 | **0.726** | **0.673** | 0.312 | **0.810** | **0.707** | 0.096 | 0.854 | **0.712** |
| Fpocket | 0.476 | 0.659 | 0.365 | 0.217 | 0.787 | 0.204 | 0.666 | 0.669 | 0.468 | 0.239 | 0.787 | 0.218 | 0.059 | 0.908 | 0.070 |
| SiteHound | 0.437 | 0.585 | 0.314 | 0.201 | 0.811 | 0.340 | 0.608 | 0.618 | 0.434 | 0.223 | 0.825 | 0.349 | 0.012 | **0.995** | 0.336 |
| MetaPocket 2.0 | 0.513 | 0.716 | 0.380 | **0.420** | 0.752 | 0.339 | 0.736 | 0.716 | 0.540 | **0.466** | 0.752 | 0.396 | 0.030 | 0.934 | 0.465 |
| DeepSite | 0.465 | 0.639 | 0.366 | 0.210 | 0.725 | 0.355 | 0.647 | 0.639 | 0.473 | 0.227 | 0.725 | 0.357 | **0.175** | 0.731 | 0.355 |
| P2Rank | 0.589 | 0.813 | 0.511 | 0.232 | 0.892 | 0.516 | 0.821 | 0.820 | 0.706 | 0.252 | 0.895 | 0.520 | 0.105 | 0.915 | 0.516 |
| BiteNet | **0.599** | 0.826 | 0.547 | 0.252 | 0.915 | 0.576 | **0.836** | 0.836 | 0.781 | 0.275 | 0.918 | 0.578 | 0.152 | 0.944 | 0.583 |
| BiteNet+ | 0.598 | **0.830** | **0.587** | 0.227 | **0.941** | **0.633** | 0.833 | **0.839** | **0.789** | 0.253 | **0.951** | **0.638** | 0.060 | 0.987 | **0.639** |
| Fpocket | 0.269 | 0.370 | 0.119 | 0.111 | 0.456 | 0.060 | 0.374 | 0.374 | 0.149 | 0.122 | 0.456 | 0.064 | 0.027 | 0.534 | 0.018 |
| SiteHound | 0.296 | 0.396 | 0.142 | 0.146 | 0.585 | 0.170 | 0.415 | 0.415 | 0.196 | 0.160 | 0.590 | 0.174 | 0.010 | 0.788 | 0.174 |
| MetaPocket 2.0 | 0.278 | 0.383 | 0.102 | **0.230** | 0.525 | 0.120 | 0.399 | 0.389 | 0.149 | **0.257** | 0.528 | 0.145 | 0.026 | 0.799 | 0.139 |
| DeepSite | 0.245 | 0.334 | 0.111 | 0.111 | 0.384 | 0.114 | 0.337 | 0.334 | 0.138 | 0.119 | 0.384 | 0.114 | 0.093 | 0.387 | 0.114 |
| P2Rank | 0.377 | 0.515 | 0.197 | 0.147 | 0.557 | 0.196 | 0.525 | 0.518 | 0.272 | 0.159 | 0.557 | 0.197 | 0.067 | 0.574 | 0.197 |
| BiteNet | 0.506 | 0.698 | 0.416 | 0.213 | 0.774 | 0.438 | 0.701 | 0.702 | 0.588 | 0.232 | 0.774 | 0.439 | **0.127** | 0.793 | 0.441 |
| BiteNet+ | **0.507** | **0.705** | **0.463** | 0.197 | **0.816** | **0.503** | **0.708** | **0.715** | **0.622** | 0.219 | **0.823** | **0.506** | 0.053 | **0.862** | **0.507** |
| Performance on HOLO4K dataset based on all protein entries | | | | | | | | | | | | | | | |
| Fpocket | 0.624 | 0.287 | 0.180 | 0.401 | 0.514 | 0.206 | 0.491 | 0.526 | 0.262 | 0.345 | 0.632 | 0.219 | 0.082 | 0.784 | 0.065 |
| SiteHound | 0.561 | 0.273 | 0.143 | 0.338 | 0.493 | 0.217 | 0.435 | 0.513 | 0.235 | 0.301 | 0.638 | 0.258 | 0.021 | **0.973** | 0.289 |
| MetaPocket 2.0 | 0.648 | 0.379 | 0.217 | 0.509 | 0.431 | 0.210 | 0.536 | 0.429 | 0.208 | 0.468 | 0.503 | 0.225 | 0.065 | 0.801 | 0.384 |
| DeepSite | 0.695 | 0.326 | 0.234 | 0.430 | 0.482 | 0.311 | 0.621 | 0.470 | 0.327 | 0.419 | 0.490 | 0.316 | **0.376** | 0.491 | 0.316 |
| P2Rank | 0.818 | 0.397 | 0.310 | 0.461 | 0.646 | 0.497 | 0.612 | 0.695 | 0.543 | 0.372 | 0.748 | 0.558 | 0.132 | 0.793 | 0.563 |
| BiteNet | 0.912 | **0.418** | 0.400 | **0.572** | 0.701 | 0.656 | **0.759** | 0.791 | 0.747 | **0.484** | 0.838 | 0.765 | 0.294 | 0.864 | 0.773 |
| BiteNet+ | **0.920** | 0.416 | **0.402** | 0.524 | **0.706** | **0.667** | 0.753 | **0.808** | **0.770** | 0.438 | **0.856** | **0.785** | 0.141 | 0.900 | **0.792** |
| Fpocket | 0.413 | 0.329 | 0.136 | 0.264 | 0.585 | 0.152 | 0.521 | 0.536 | 0.281 | 0.282 | 0.681 | 0.191 | 0.053 | 0.857 | 0.045 |
| SiteHound | 0.391 | 0.317 | 0.120 | 0.233 | 0.562 | 0.174 | 0.481 | 0.509 | 0.233 | 0.256 | 0.681 | 0.214 | 0.012 | **0.999** | 0.221 |
| MetaPocket 2.0 | 0.475 | 0.458 | 0.188 | 0.369 | 0.510 | 0.178 | 0.592 | 0.484 | 0.257 | **0.403** | 0.549 | 0.217 | 0.043 | 0.892 | 0.317 |
| DeepSite | 0.481 | 0.382 | 0.191 | 0.289 | 0.545 | 0.241 | 0.650 | 0.523 | 0.361 | 0.309 | 0.554 | 0.252 | **0.252** | 0.556 | 0.245 |
| P2Rank | 0.579 | 0.471 | 0.267 | 0.315 | 0.744 | 0.411 | 0.726 | 0.767 | 0.603 | 0.319 | 0.832 | 0.473 | 0.086 | 0.874 | 0.451 |
| BiteNet | 0.637 | **0.502** | 0.364 | **0.386** | 0.819 | 0.582 | 0.849 | 0.872 | 0.817 | 0.396 | 0.941 | 0.672 | 0.190 | 0.971 | 0.664 |
| BiteNet+ | **0.639** | **0.502** | **0.381** | 0.352 | **0.824** | **0.617** | **0.861** | **0.881** | **0.844** | 0.367 | **0.947** | **0.715** | 0.088 | 0.986 | **0.702** |
| Fpocket | 0.226 | 0.174 | 0.039 | 0.134 | 0.309 | 0.040 | 0.276 | 0.278 | 0.076 | 0.142 | 0.361 | 0.050 | 0.022 | 0.462 | 0.010 |
| SiteHound | 0.260 | 0.203 | 0.047 | 0.161 | 0.376 | 0.074 | 0.321 | 0.323 | 0.094 | 0.175 | 0.448 | 0.089 | 0.010 | 0.767 | 0.100 |
| MetaPocket 2.0 | 0.256 | 0.235 | 0.051 | 0.206 | 0.329 | 0.059 | 0.296 | 0.261 | 0.063 | 0.218 | 0.352 | 0.069 | 0.035 | 0.694 | 0.101 |
| DeepSite | 0.272 | 0.209 | 0.062 | 0.158 | 0.290 | 0.072 | 0.355 | 0.275 | 0.107 | 0.169 | 0.293 | 0.075 | 0.137 | 0.294 | 0.073 |
| P2Rank | 0.357 | 0.276 | 0.081 | 0.190 | 0.427 | 0.117 | 0.438 | 0.442 | 0.175 | 0.193 | 0.483 | 0.138 | 0.052 | 0.510 | 0.131 |
| BiteNet | 0.565 | 0.435 | 0.295 | **0.341** | 0.705 | 0.469 | 0.743 | 0.742 | 0.646 | **0.348** | 0.803 | 0.534 | **0.167** | 0.831 | 0.529 |
| BiteNet+ | **0.572** | **0.441** | **0.317** | 0.317 | **0.726** | **0.518** | **0.760** | **0.762** | **0.691** | 0.325 | **0.823** | **0.590** | 0.079 | **0.859** | **0.579** |
| Performance on COACH420 dataset based on all protein entries for which there are at least one relevant ligand exists respectively to the true positive criteria. | | | | | | | | | | | | | | | |
| Fpocket | 0.648 | 0.490 | 0.339 | 0.315 | 0.634 | 0.215 | 0.519 | 0.555 | 0.321 | 0.292 | 0.663 | 0.211 | 0.086 | 0.755 | 0.074 |
| SiteHound | 0.585 | 0.439 | 0.289 | 0.285 | 0.641 | 0.329 | 0.469 | 0.531 | 0.329 | 0.257 | 0.668 | 0.333 | 0.020 | **0.933** | 0.338 |
| MetaPocket 2.0 | 0.714 | 0.556 | 0.393 | **0.591** | 0.602 | 0.373 | 0.656 | 0.587 | 0.386 | **0.559** | 0.625 | 0.373 | 0.048 | 0.811 | 0.510 |
| DeepSite | 0.658 | 0.486 | 0.372 | 0.308 | 0.572 | 0.377 | 0.540 | 0.517 | 0.387 | 0.295 | 0.575 | 0.378 | **0.256** | 0.577 | 0.378 |
| P2Rank | 0.810 | 0.623 | 0.536 | 0.332 | 0.707 | 0.552 | 0.642 | 0.666 | 0.557 | 0.309 | 0.723 | 0.557 | 0.152 | 0.740 | 0.553 |
| BiteNet | 0.855 | **0.653** | **0.612** | 0.380 | 0.769 | 0.679 | **0.688** | 0.722 | 0.661 | 0.360 | 0.789 | 0.687 | 0.232 | 0.809 | 0.691 |
| BiteNet+ | **0.861** | 0.651 | 0.612 | 0.343 | **0.777** | **0.693** | 0.663 | **0.730** | **0.674** | 0.311 | **0.807** | **0.704** | 0.094 | 0.852 | **0.709** |
| Fpocket | 0.690 | 0.659 | 0.473 | 0.314 | 0.787 | 0.264 | 0.666 | 0.669 | 0.468 | 0.310 | 0.787 | 0.261 | 0.085 | 0.908 | 0.089 |
| SiteHound | 0.620 | 0.585 | 0.408 | 0.285 | 0.811 | 0.439 | 0.608 | 0.618 | 0.434 | 0.284 | 0.825 | 0.443 | 0.017 | **0.995** | 0.435 |
| MetaPocket 2.0 | 0.743 | 0.716 | 0.553 | **0.609** | 0.752 | 0.481 | 0.736 | 0.716 | 0.540 | **0.603** | 0.752 | 0.479 | 0.043 | 0.934 | 0.654 |
| DeepSite | 0.669 | 0.639 | 0.474 | 0.299 | 0.725 | 0.470 | 0.647 | 0.639 | 0.473 | 0.298 | 0.725 | 0.470 | **0.248** | 0.731 | 0.470 |
| P2Rank | 0.851 | 0.813 | 0.705 | 0.330 | 0.892 | 0.711 | 0.821 | 0.820 | 0.706 | 0.327 | 0.895 | 0.713 | 0.151 | 0.915 | 0.712 |
| BiteNet | 0.866 | 0.821 | 0.772 | 0.368 | 0.913 | 0.823 | 0.829 | 0.833 | 0.777 | 0.363 | 0.917 | 0.824 | 0.227 | 0.948 | 0.836 |
| BiteNet+ | **0.874** | **0.833** | **0.780** | 0.329 | **0.933** | **0.844** | **0.833** | **0.841** | **0.787** | 0.327 | **0.944** | **0.849** | 0.090 | 0.984 | **0.856** |
| Fpocket | 0.390 | 0.370 | 0.153 | 0.160 | 0.456 | 0.077 | 0.374 | 0.374 | 0.149 | 0.158 | 0.456 | 0.076 | 0.038 | 0.534 | 0.023 |
| SiteHound | 0.420 | 0.396 | 0.183 | 0.207 | 0.585 | 0.218 | 0.415 | 0.415 | 0.196 | 0.204 | 0.590 | 0.219 | 0.014 | 0.788 | 0.224 |
| MetaPocket 2.0 | 0.403 | 0.383 | 0.149 | **0.333** | 0.525 | 0.170 | 0.399 | 0.389 | 0.149 | **0.332** | 0.528 | 0.175 | 0.037 | 0.799 | 0.196 |
| DeepSite | 0.352 | 0.334 | 0.141 | 0.157 | 0.384 | 0.148 | 0.337 | 0.334 | 0.138 | 0.157 | 0.384 | 0.148 | 0.133 | 0.387 | 0.148 |
| P2Rank | 0.545 | 0.515 | 0.272 | 0.209 | 0.557 | 0.270 | 0.525 | 0.518 | 0.272 | 0.206 | 0.557 | 0.270 | 0.096 | 0.574 | 0.271 |
| BiteNet | 0.739 | 0.702 | 0.596 | 0.315 | 0.782 | 0.633 | 0.702 | 0.706 | 0.594 | 0.310 | 0.782 | 0.633 | **0.193** | 0.806 | 0.639 |
| BiteNet+ | **0.744** | **0.710** | **0.613** | 0.282 | **0.802** | **0.664** | **0.710** | **0.718** | **0.619** | 0.280 | **0.810** | **0.667** | 0.077 | **0.849** | **0.672** |
| Performance on HOLO4K dataset based on all protein entries for which there are at least one relevant ligand exists respectively to the true positive criteria. | | | | | | | | | | | | | | | |
| Fpocket | 0.624 | 0.287 | 0.180 | 0.401 | 0.514 | 0.206 | 0.491 | 0.526 | 0.262 | 0.345 | 0.632 | 0.219 | 0.082 | 0.784 | 0.065 |
| SiteHound | 0.561 | 0.273 | 0.143 | 0.338 | 0.493 | 0.217 | 0.435 | 0.513 | 0.235 | 0.301 | 0.638 | 0.258 | 0.021 | **0.973** | 0.289 |
| MetaPocket 2.0 | 0.648 | 0.379 | 0.217 | 0.509 | 0.431 | 0.210 | 0.536 | 0.429 | 0.208 | 0.468 | 0.503 | 0.225 | 0.065 | 0.801 | 0.384 |
| DeepSite | 0.695 | 0.326 | 0.234 | 0.430 | 0.482 | 0.311 | 0.621 | 0.470 | 0.327 | 0.419 | 0.490 | 0.316 | **0.376** | 0.491 | 0.316 |
| P2Rank | 0.818 | 0.397 | 0.310 | 0.461 | 0.646 | 0.497 | 0.612 | 0.695 | 0.543 | 0.372 | 0.748 | 0.558 | 0.132 | 0.793 | 0.563 |
| BiteNet | 0.913 | **0.424** | 0.407 | **0.566** | 0.702 | 0.657 | **0.757** | 0.791 | 0.748 | **0.481** | 0.838 | 0.765 | 0.294 | 0.864 | 0.773 |
| BiteNet+ | **0.919** | 0.422 | **0.408** | 0.517 | **0.708** | **0.667** | 0.749 | **0.808** | **0.770** | 0.433 | **0.856** | **0.784** | 0.140 | 0.900 | **0.791** |
| Fpocket | 0.584 | 0.329 | 0.194 | 0.373 | 0.585 | 0.218 | 0.521 | 0.536 | 0.281 | 0.343 | 0.681 | 0.234 | 0.075 | 0.857 | 0.064 |
| SiteHound | 0.553 | 0.317 | 0.162 | 0.329 | 0.562 | 0.236 | 0.481 | 0.509 | 0.233 | 0.311 | 0.681 | 0.268 | 0.017 | **0.999** | 0.302 |
| MetaPocket 2.0 | 0.662 | 0.458 | 0.270 | 0.514 | 0.510 | 0.249 | 0.592 | 0.484 | 0.257 | **0.493** | 0.549 | 0.254 | 0.060 | 0.892 | 0.439 |
| DeepSite | 0.680 | 0.382 | 0.269 | 0.409 | 0.545 | 0.341 | 0.650 | 0.523 | 0.361 | 0.406 | 0.554 | 0.347 | **0.356** | 0.556 | 0.347 |
| P2Rank | 0.821 | 0.471 | 0.366 | 0.445 | 0.744 | 0.564 | 0.726 | 0.767 | 0.603 | 0.389 | 0.832 | 0.619 | 0.121 | 0.874 | 0.622 |
| BiteNet | 0.907 | **0.513** | 0.488 | **0.544** | 0.823 | 0.760 | 0.848 | 0.873 | 0.821 | 0.486 | 0.942 | 0.853 | 0.272 | 0.971 | 0.866 |
| BiteNet+ | **0.911** | 0.512 | **0.494** | 0.492 | **0.826** | **0.777** | **0.859** | **0.883** | **0.847** | 0.441 | **0.946** | **0.871** | 0.125 | 0.987 | **0.885** |
| Fpocket | 0.318 | 0.174 | 0.055 | 0.189 | 0.309 | 0.057 | 0.275 | 0.278 | 0.076 | 0.172 | 0.360 | 0.061 | 0.031 | 0.462 | 0.014 |
| SiteHound | 0.367 | 0.203 | 0.064 | 0.228 | 0.376 | 0.100 | 0.322 | 0.323 | 0.094 | 0.213 | 0.448 | 0.111 | 0.014 | 0.767 | 0.136 |
| MetaPocket 2.0 | 0.354 | 0.234 | 0.072 | 0.286 | 0.328 | 0.082 | 0.295 | 0.260 | 0.063 | 0.266 | 0.352 | 0.081 | 0.048 | 0.693 | 0.138 |
| DeepSite | 0.385 | 0.209 | 0.088 | 0.224 | 0.290 | 0.102 | 0.355 | 0.275 | 0.107 | 0.221 | 0.293 | 0.103 | 0.194 | 0.294 | 0.103 |
| P2Rank | 0.505 | 0.276 | 0.111 | 0.268 | 0.426 | 0.161 | 0.438 | 0.442 | 0.175 | 0.236 | 0.482 | 0.180 | 0.073 | 0.510 | 0.182 |
| BiteNet | 0.808 | 0.446 | 0.395 | **0.484** | 0.712 | 0.613 | 0.748 | 0.747 | 0.653 | **0.429** | 0.809 | 0.680 | **0.241** | 0.836 | 0.690 |
| BiteNet+ | **0.813** | **0.449** | **0.406** | 0.444 | **0.731** | **0.650** | **0.763** | **0.766** | **0.693** | 0.393 | **0.826** | **0.717** | 0.111 | **0.861** | **0.724** |

Table 2: List of the PDB identifiers filtered on the dataset preprocessing stage

| Type | Pdb code |
|---|---|
| unknown ligands | UNL |
| detergents | 12P, 15P, 1PE, 2CV, 2PE, 78M, 78N, BCR, BNG, BOG, BTB, C14, C8E, CDL, CLR, CM5, CPS, DAO, DGA, DGD, DMU, DXC, L2P, LDA, LFA, LHG, LI1, LMG, LMT, LMU, MPG, MQ7, MTN, MYR, MYS, NS5, OLA, OLB, OLC, P4C, P6G, PC1, PCW, PE4, PE5, PE8, PEE, PG6, PGV, PL9, PLC, OLM, PTY, PX4, SPN, SPO, SQD, STE, TGL, U10, YO1, HOH |
| modified residues | CSD, HYP, BMT, 5HP, ABA, AIB, CSW, OCS, DAL, DAR, DSG, DSP, DCY, CRO, DGL, DGN, DHI, DIL, DIV, DLE, DLY, DPN, DPR, DSN, DTH, DTR, DTY, DVA, CGU, KCX, LLP, CXM, FME, MLE, MVA, NLE, PTR, ORN, SEP, TPO, PCA, PVL, SAR, CEA, CSO, CSS, CSX, CME, TYS, TPQ, STY, NH2, CBX, ACE, FOR, IVA, BOC, LYR, MSE |
| cofactors | ADP, AMP, ATP, CMP, COA, FAD, FMN, NAP, NDP |
| carbohydrates | BGC, GLC, MAN, BMA, FUC, GAL, GLA, NAG, NGA, SIA, XYS |

Source: `https://www.globalphasing.com/buster/manual/maketnt/manual/lib_val/library_validation.html`

Table 3: Performance of the deep learning models trained with different parameters.

| Voxel size, Å | Density cutoff, Å | Cube size | Cell size | Stride | $\gamma$ | $\lambda$ | Minibatch | Average precision |
|---|---|---|---|---|---|---|---|---|
| 1.0 | 2.0 | 48 | 4 | 32 | 1e-05 | 5 | 64 | 0.307 |
| 1.0 | 2.0 | 48 | 8 | 32 | 1e-05 | 5 | 64 | 0.394 |
| 1.0 | 2.0 | 48 | 16 | 32 | 1e-05 | 5 | 64 | 0.353 |
| 1.0 | 2.0 | 64 | 16 | 32 | 1e-05 | 5 | 32 | 0.383 |
| 1.0 | 2.0 | 64 | 4 | 32 | 1e-05 | 5 | 32 | 0.314 |
| 1.0 | 2.0 | 64 | 8 | 32 | 1e-05 | 5 | 4 | 0.387 |
| 1.0 | 2.0 | 64 | 8 | 32 | 1e-05 | 5 | 8 | 0.331 |
| 1.0 | 2.0 | 64 | 8 | 32 | 1e-05 | 5 | 16 | 0.429 |
| 1.0 | 2.0 | 64 | 8 | 32 | 1e-05 | 5 | 32 | 0.404 |
| 1.0 | 1.0 | 80 | 16 | 32 | 1e-05 | 5 | 16 | 0.365 |
| 1.0 | 2.0 | 80 | 16 | 32 | 1e-05 | 5 | 16 | 0.343 |
| 1.0 | 4.0 | 80 | 16 | 32 | 1e-05 | 5 | 16 | 0.387 |
| 1.0 | 8.0 | 80 | 16 | 32 | 1e-05 | 5 | 16 | 0.355 |
| 1.0 | 16.0 | 80 | 16 | 32 | 1e-05 | 5 | 16 | 0.368 |
| 1.0 | 2.0 | 80 | 16 | 32 | 0 | 5 | 16 | 0.372 |
| 1.0 | 2.0 | 80 | 16 | 32 | 1e-01 | 5 | 16 | 0.216 |
| 1.0 | 2.0 | 80 | 16 | 32 | 1e-02 | 5 | 16 | 0.264 |
| 1.0 | 2.0 | 80 | 16 | 32 | 1e-03 | 5 | 16 | 0.299 |
| 1.0 | 2.0 | 80 | 16 | 32 | 1e-04 | 5 | 16 | 0.362 |
| 1.0 | 2.0 | 80 | 16 | 32 | 1e-05 | 5 | 16 | 0.371 |
| 1.0 | 2.0 | 80 | 16 | 32 | 1e-06 | 5 | 16 | 0.351 |
| 1.0 | 2.0 | 80 | 16 | 32 | 1e-05 | 1 | 16 | 0.368 |
| 1.0 | 2.0 | 80 | 16 | 32 | 1e-05 | 10 | 16 | 0.377 |
| 1.0 | 2.0 | 80 | 16 | 32 | 1e-05 | 100 | 16 | 0.333 |
| 1.0 | 2.0 | 80 | 16 | 32 | 1e-05 | 1000 | 16 | 0.346 |
| 1.0 | 2.0 | 80 | 16 | 16 | 1e-05 | 5 | 16 | 0.272 |
| 1.0 | 2.0 | 80 | 16 | 24 | 1e-05 | 5 | 16 | 0.354 |
| 1.0 | 2.0 | 80 | 16 | 32 | 1e-05 | 5 | 16 | 0.399 |
| 1.0 | 2.0 | 80 | 16 | 48 | 1e-05 | 5 | 16 | 0.400 |
| 1.0 | 2.0 | 80 | 4 | 32 | 1e-05 | 5 | 16 | 0.302 |
| 1.0 | 2.0 | 80 | 8 | 32 | 1e-05 | 5 | 16 | 0.387 |
| 0.8 | 1.6 | 64 | 8 | 32 | 1e-05 | 5 | 32 | 0.435 |
| 1.5 | 3.0 | 64 | 8 | 32 | 1e-05 | 5 | 32 | 0.332 |
| 2.0 | 4.0 | 64 | 8 | 32 | 1e-05 | 5 | 32 | 0.283 |
| 3.0 | 6.0 | 64 | 8 | 32 | 1e-05 | 5 | 32 | 0.147 |

Table 4: Number of proteins in datasets. For each method only subset of proteins for which there were successful prediction and that were not present in the training set were considered.

| | COACH420 | HOLO4K |
|---|---|---|
| | 420 | 4542 |
| Fpocket | 420 | 4009 |
| SiteHound | 420 | 2878 |
| MetaPocket 2.0 | 417 | 2575 |
| DeepSite | 420 | 3991 |
| P2Rank | 420 | 4542 |
| BiteNet | 363 | 4186 |

Table 5: Average number of ligands and predictions for datasets. The first row (ligands_P2Rank) is average number of ligands chosen as in P2Rank paper. The second row (ligands_BiteNet)is for ligands considered relevant as in our method. The rest corresponds to the average number of predicted binding sites. For BiteNet the listed values correspond to the probability scores of 0.01 and 0.1, respectively, used in the benchmarks.

| | COACH420 | HOLO4K |
|---|---|---|
| ligands_P2Rank | 1.39 | 2.36 |
| ligands_BiteNet | 0.73 | 1.30 |
| Fpocket | 14.60 | 27.01 |
| SiteHound | 60.23 | 99.87 |
| MetaPocket 2.0 | 22.56 | 22.74 |
| DeepSite | 3.16 | 2.79 |
| P2Rank | 6.26 | 12.57 |
| BiteNet | 4.61/1.52 | 6.35/2.57 |

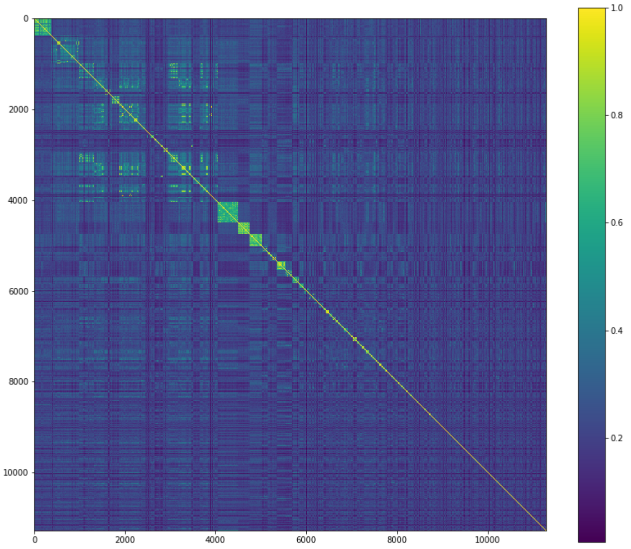


Figure 4: Pair-wise structure similarity matrix calculated for the protein chains in the training dataset.
